# Genetic underpinnings of predicted changes in cardiovascular function using self supervised learning

**DOI:** 10.1101/2024.08.15.608061

**Authors:** Zachary Levine, Guy Lutsker, Anastasia Godneva, Adina Weinberger, Maya Pompan, Yeela Talmor-Barkan, Yotam Reisner, Hagai Rossman, Eran Segal

## Abstract

**Background:** The genetic underpinnings of cardiovascular disease remain elusive. Contrastive learning algorithms have recently shown cutting-edge performance in extracting representations from electrocardiogram (ECG) signals that characterize cross-temporal cardiovascular state. However, there is currently no connection between these representations and genetics.

**Methods:** We designed a new metric, denoted as Delta ECG, which measures temporal shifts in patients’ cardiovascular state, and inherently adjusts for inter-patient differences at baseline. We extracted this measure for 4,782 patients in the Human Phenotype Project using a novel self-supervised learning model, and quantified the associated genetic signals with Genome-Wide-Association Studies (GWAS). We predicted the expression of thousands of genes extracted from Peripheral Blood Mononuclear Cells (PBMCs). Downstream, we ran enrichment and overrepresentation analysis of genes we identified as significantly predicted from ECG.

**Findings:** In a Genome-Wide Association Study (GWAS) of Delta ECG, we identified five associations that achieved genome-wide significance. From baseline embeddings, our models significantly predict the expression of 57 genes in men and 9 in women. Enrichment analysis showed that these genes were predominantly associated with the electron transport chain and the same immune pathways as identified in our GWAS.

**Conclusions:** We validate a novel method integrating self-supervised learning in the medical domain and simple linear models in genetics. Our results indicate that the processes underlying temporal changes in cardiovascular health share a genetic basis with CVD, its major risk factors, and its known correlates. Moreover, our functional analysis confirms the importance of leukocytes, specifically eosinophils and mast cells with respect to cardiac structure and function.

## Introduction

Electrocardiography (ECG) is a critical tool in diagnostic cardiology, and exploring the genetic underpinnings of CVD though it remains an open but promising problem^1^. Existing works that apply deep learning to ECG either explore genetics but fix a supervisory signal in the form of target labels for age^2^ and Atrial Fibrillation (AF)^3^, or learn embeddings using Self-Supervised-Learning (SSL), but do not consider genetic implications of changes in these features over time^4,5^. As well, these supervisory signals potentially bias learned features away from general health state, towards these individual target labels^3^. The branch of machine learning algorithms that use neural networks to learn discriminative features from data without target supervisory labels is known as SSL. These feature vectors, often denoted as embeddings, are traditionally learned through masked signal modeling or contrastive learning ^6^.

We used ECG recordings and genetics data (DNA, RNA expression) from the Human Phenotype Project (HPP) for this project. The HPP is a large-scale longitudinal study, centered on the deep phenotyping of people between 22 and 70 years of age in Israel. Over the course of several years, the study has collected a wide range of clinical data and biological data, and seeks to discover correlates for disease and targets for disease ^7^. Recordings were collected at intake appointments, and repeated two years later for ~40% of the study cohort (see Methods).

When neural networks are trained based on contrastive learning, coordinates in the latent space of the model are expected to correspond to various parameters of a patient’s health state ^4,5^. We quantified each patient’s temporal shift in cardiovascular state through the distance between their baseline and follow up ECG embeddings. We defined our new metric, Delta ECG, as the cosine similarity between the embeddings of first and second appointments for each patient. This single number measures a patient’s changes in cardiovascular performance over a two year period, relative to their intake visit. As such, it functions as a valuable measure of cardiovascular aging over time, while inherently adjusting for baseline patient health differences across the HPP population.

In our GWAS of Delta ECG, five SNPS reached genome-wide significance. Our pathway analyses and subsequent RNA expression predictions indicate that Delta ECG shares a genetic basis with CVD, its major risk factors, and its known correlates in the immune system. Our raw embeddings were directly able to predict the expression of genes along the pathways containing the GWAS hits for Delta ECG, validating both our findings and our method.

## Results

### Description of the Pooled HPP Cohort

The present study used the cohort from the Human Phenotype Project, utilizing a pooled cohort consisting of 16,775 ECG recordings from 11,933 patients (diseased and healthy), subdivided into two non overlapping four second windows each. For a complete list of exclusion criteria, we direct readers to the original paper describing the cohort^7^. This distribution, along with descriptive summary statistics, can be found in Tables 1 and 2. To understand the vast complexities of human health, including both function and dysfunction we pooled diseased and healthy individuals into one large dataset to train our model.

**Table 1:** Baseline features of included healthy cohorts, stratified by age.

| Table 1: Baseline features of Healthy Cohort |  |  |  |
| --- | --- | --- | --- |
| Numerical Features | Healthy age > 50 | Healthy age < 30 | Healthy Other |
|  | Mean (SD) | Mean (SD) | Mean (SD) |
| Age at Baseline (years) | 51.86 (8.01) | 26.57 (4.02) | 53.05 (19.09) |
| BMI | 26.07 (4.16) | 24.42 (4.08) | 26.95 (3.67) |
| Waist | 88.77 (12.04) | 78.71 (12.11) | 88.45 (12.9) |
| Height | 169.07 (9.19) | 168.79 (9.33) | 169.77 (6.67) |
| Sitting blood pressure (systolic) | 119.85 (16.6) | 111.79 (15.26) | 127.38 (14.58) |
| Sitting blood pressure (diastolic) | 77.76 (10.34) | 66.22 (9.83) | 66.31 (7.5) |

**Table 2:** Baseline features of included disease cohorts.

| Table 2: Baseline features of Diseased Cohorts |  |  |  |  |  |
| --- | --- | --- | --- | --- | --- |
| Feature | CVD | Colorectal Cancer Recovered | Breast Cancer Recovered | BRCA | Endometriosis |
|  |  | Mean (SD) | Mean (SD) | Mean (SD) | Mean (SD) |
| Age at baseline (years) | 57.11 (9.56) | 56.38 (10.33) | 49.83 (10.01) | 43.08 (10.13) | 31.34 (10.77) |
| BMI | 28.61 (4.45) | 26.96 (3.8) | 26.03 (5.2) | 25.27 (4.18) | 24.12 (4.77) |
| Waist Circumference | 98.96 (12.75) | 93.0 (14.2) | 84.1 (13.09) | 83.05 (12.31) | 76.55 (11.48) |
| Height (cm) | 171.43 (8.81) | 172.2 (8.74) | 162.68 (6.42) | 165.65 (7.91) | 162.19 (6.36) |
| Sitting blood pressure (systolic) | 128.11 (17.56) | 130.88 (28.6) | 117.45 (18.37) | 112.45 (14.05) | 105.55 (11.6) |
| Sitting blood pressure (diastolic) | 79.2 (9.97) | 73.19 (13.57) | 71.89 (11.47) | 70.79 (10.44) | 67.16 (8.65) |

### Self-Supervised Learning Objective

In current literature, there are two leading methods of conducting self-supervised contrastive learning with ECG data: Contrastive Learning of Cardiac Signals (CLOCS/CMSC) ^4^, and Patient Contrastive Learning (PCLR) ^5^. To capitalize on the strengths of our dataset, a high number of repeat visits and high temporal resolution, we trained a model using a combination of both approaches. Overall, the training objective of the model, given two different views of an ECG signal, was to maximize the agreement between learned representations if those two windows are taken from the same patient, and to minimize them otherwise. Our complete training and validation curves can be found in Figure 2, and architecture specifications/hyperparameters can be found in the Method section of this work.

**Figure 1:**
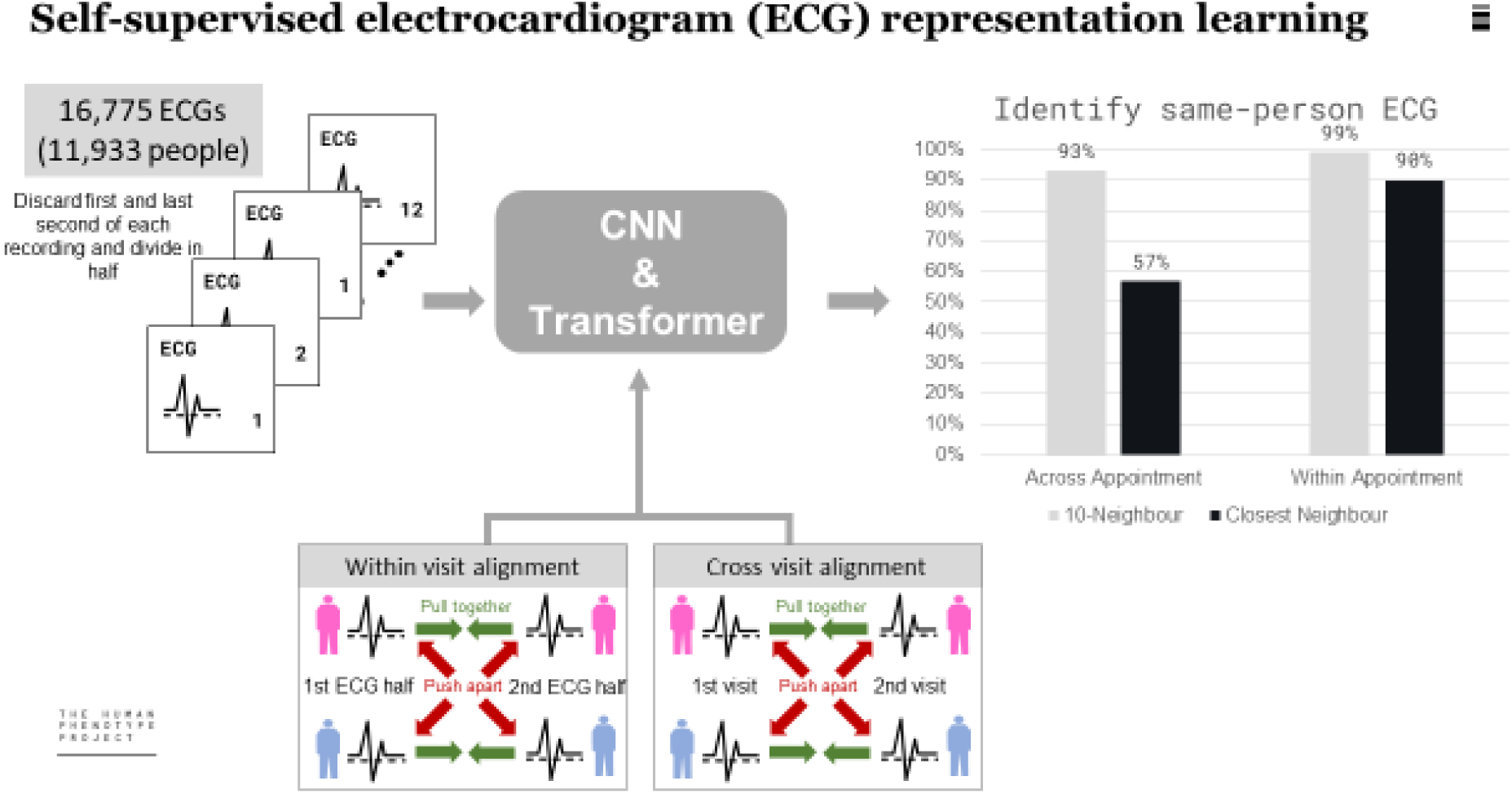
Our model correctly matches patients to their corresponding ECG recordings in the latent space, both across appointments and within them. We split ECG recordings in time, and passed them through our CNN/Transformer network. We trained our model based on a combination of the two leading contrastive learning algorithms for ECGs: Contrastive Multi-Segment Coding (bottom, left) and Patient Contrastive Learning (bottom, right). Our patient identification performance on these two tasks is shown in the associated barplot.

**Figure 2:**
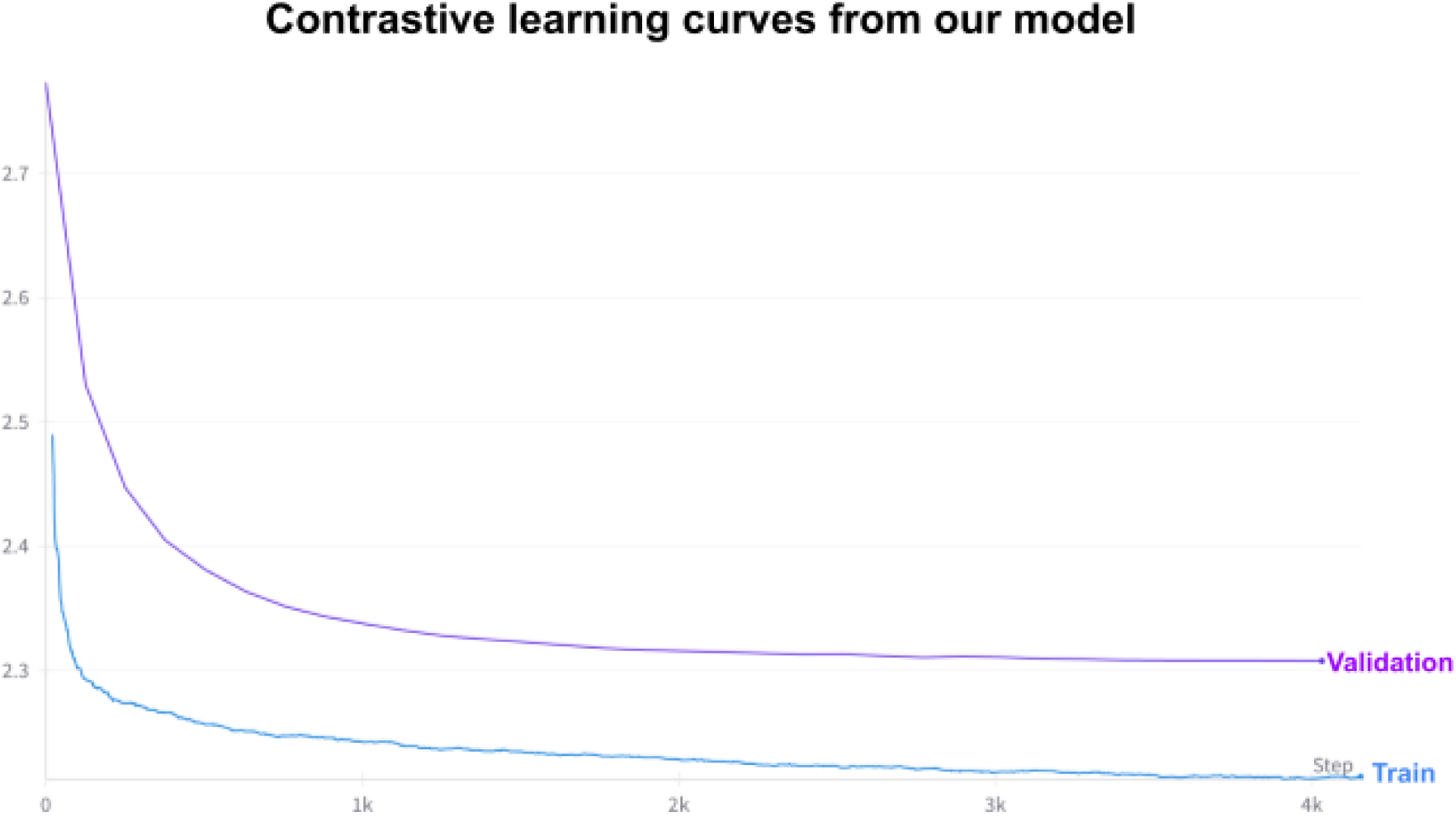
The model converges based on contrastive loss after many iterations. a) Training loss (blue) and validation loss (purple) of the trained model. We initialized the model (Table S1, See Supplement) with random weights, and trained using the AdamW, with an initial learning rate of 1e-5. We reduced the learning rate by a factor of 5 for every 5 epochs that the validation loss did not decrease, and after 15 epochs of non-decreasing validation loss, we stopped training the model, and checkpointed the weights for later use.

### Model Validation: Patient Identification

We trained our neural network (Table S1, See Supplement) until satisfaction of the stopping criteria (see Methods). To validate our model, we applied metrics over our embedding space that checked for how many patients, the closest embedding vector in the latent space to their first appointment was their second one (Top-1 across). We did the same for the matching of first and second halves of each recording (Top-1 within)^8^. Top-10 accuracy (in either case) was a relaxation so that the correct embedding need only be among the top 10 closest vectors. Our trained model achieved (test) accuracies of 57% (top-1) and 93% (top-10) across appointments. Within appointments, we achieved 90% (top-1) and 99%, (top-10). One might wonder how this performs relative to random assignment: 57% “across” patient identification is 150 times better than random. These results indicate that our learned representations effectively characterize patient health, and serve as validation for our model.

### The HPP Delta ECG distribution is as expected

We extracted two distributions: the Delta ECG for each participant and the null distribution, which was the distance between all first and all second appointments, regardless of patient identity. These two distributions are shown in Figure 3. We observed that the Delta ECG distribution is significantly to the right of the null one, which is centered around zero. This indicates that embeddings from a patient’s first appointment are much closer to embeddings from that patient’s second appointment than those from all other people.

**Figure 3:**
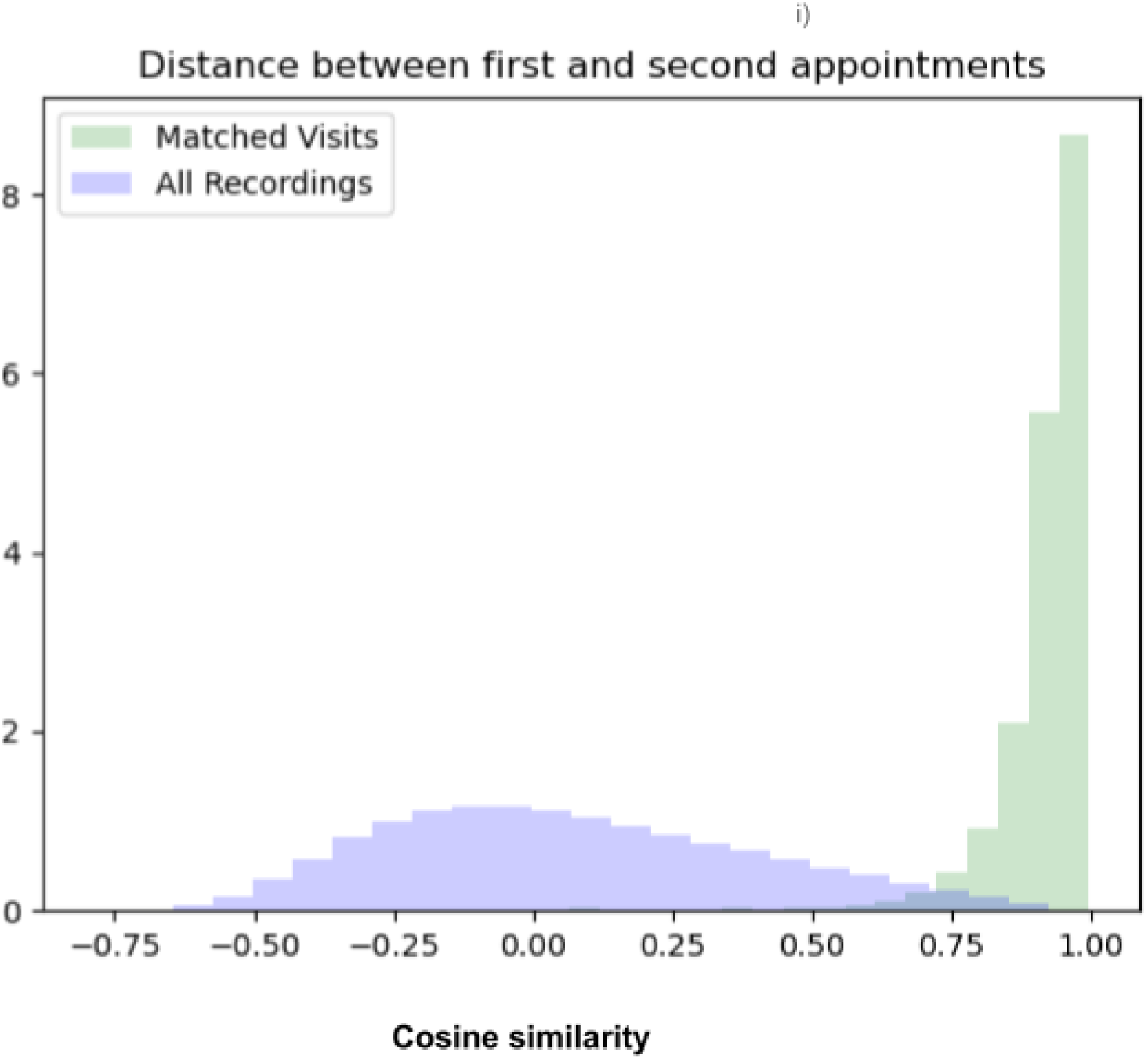
Patients are correctly matched to their repeat visits in the latent space. Distribution (density) of cosine similarity between first and second visits of matched (green) and all (blue) appointments [the null distribution].

### GWAS of Delta ECG: Five SNPs reach genome-wide significance

Starting from the DNA level, we sought to understand the genetic associations for our embeddings. We adjusted for age, gender, and the top 10 Principal Components (PCs) of the variance-standardized relationship matrix, and pruned for first degree relations. Complete methods describing GWAS protocols and our thresholds for the HPP have been detailed previously^9^. There were no multi-trait hits in GWAS of the top 10 principal components from either first or second appointment embeddings. However, we identified five genome-wide significant variants associated with Delta ECG (see Introduction) (Figure 4 and Table 3). We clumped the full unbiased GWAS results for Linkage Disequilibrium (LD), forming 629 clumps from the 1,806 top variants. Most importantly, all five of these SNPs were put in different clumps.

**Figure 4:**
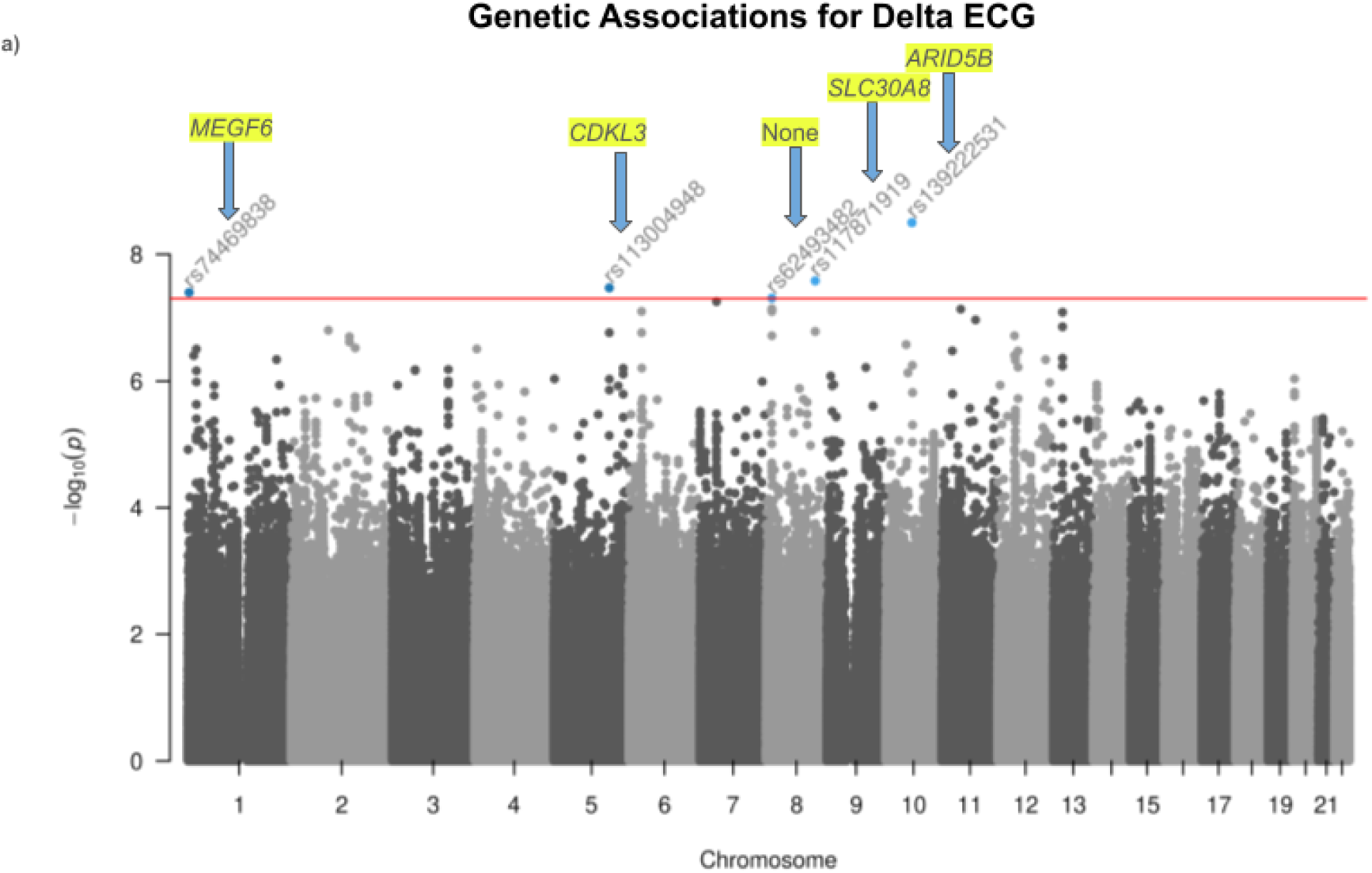
Changes in ECG cardiovascular state share a genetic basis with CVD, its known correlates, and its major risk factors. Manhattan plot of Delta ECG showing the top SNP associations for this trait, as assessed by our GWAS. Names of genes are labeled above each SNP they contain, and are highlighted in yellow.

**Table 3:**
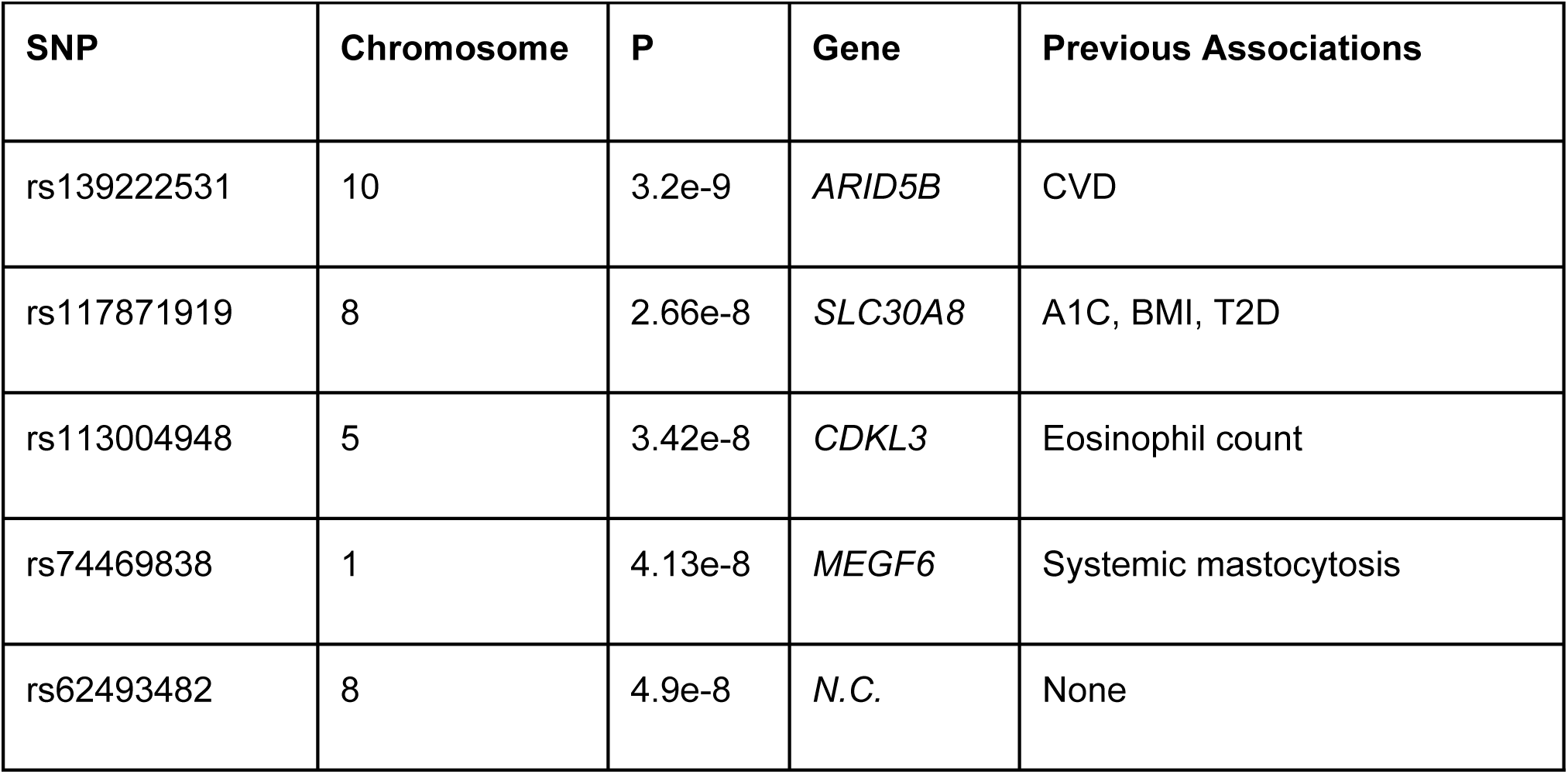
Delta ECG shares a genetic basis with CVD and its known risk factors and correlates. Significant hits for Delta ECG and their previous genetic associations.

| Table 3: Significant Hits for Delta ECG |  |  |  |  |
| --- | --- | --- | --- | --- |
| SNP | Chromosome | P | Gene | Previous Associations |
| rs139222531 | 10 | 3.2e-9 | *ARID5B* | CVD |
| rs117871919 | 8 | 2.66e-8 | *SLC30A8* | A1C, BMI, T2D |
| rs113004948 | 5 | 3.42e-8 | *CDKL3* | Eosinophil count |
| rs74469838 | 1 | 4.13e-8 | *MEGF6* | Systemic mastocytosis |
| rs62493482 | 8 | 4.9e-8 | *N.C.* | None |

### Delta ECG shares genetic underpinnings with CVD through *ARID5B*

With respect to our GWAS results, no SNP had been previously reported in any GWAS. We thus performed our analyses on the gene level. Our strongest hit (rs139222531, P < 4e-9) is in *ARID5B*, a gene with previously established significant hits for CVD within the UK Biobank^10^. *ARID5B* encodes a protein that belongs to the AT-rich interaction domain (ARID) family. Previous studies have identified this gene as playing a role in the regulation of inflammation and immune responses with respect to atherosclerosis^11^. The identification of a previously established significant hit for CVD within our framework validates the usefulness of our model.

### Common genetic architecture between Delta ECG and glycemic CVD risk factors through *SLC30A8*

However, beyond CVD itself, we identified a significant hit in *SLC30A8*, (rs117871919, P < 3e-8). *SLC30A8* encodes a zinc transporter protein that is primarily expressed in pancreatic beta cells. *SLC3A03* has known genetic associations for glycemic variance and control, namely HbA1c ^12^, BMI^12^, and T2D^13^, which is a well-established risk factor for CVD^14^. These findings indicate that Delta ECG shares a genetic signal not just with CVD, but with its significant risk factors as well.

### Delta ECG also shares a genetic basis with known correlates of CVD through *CDKL3* and *MEGF6*

We found a hit (rs113004948, P < 4e-8) in *CDKL3*, a gene that encodes a protein belonging to the cyclin-dependent kinase (CDK) family. *CDKL3 has* previously established significant hits for Eosinophil counts^15^, which are known to have a strong relationship with frequency of cardiac complications (Bozkuş et al.). As well, our weakest but still genome-wide significant hit (rs74469838, P < 5e-8) is in *MEGF6*, which has existing strong associations for Systemic Mastocytosis^16^, another correlate of CVD^17^. These results indicate that Delta ECG shares a genetic basis with correlates of CVD.

### We could predict the expression of 66 genes significantly from our model

We wanted to understand the functional predictive power of our embeddings on a cellular level. In the HPP, RNA expression levels are collected from a sample containing a bulk of peripheral blood mononuclear cells (PBMCs) in a process known as RNA-Sequencing, or RNASeq (see Methods). After correction, in men, we were able to predict 57 genes significantly at a threshold of P<0.05, and 9 genes at the more stringent (0.01) level. In women, we could predict 9 genes at 0.05, and 0 at the more stringent testing threshold, after correction. These genes are displayed in Table S2. Among them are prominent mitochondrial genes such as *MT-RNR1*/*2* and *MT-TV*.

We aimed to explore which biological pathways were implicated amongst the genes whose expression values we could predict significantly in men and women. To this end, we ran overrepresentation analysis over three major gene sets: GO Biological Processes 2023, MSigDB_Hallmark_2020, and Kegg 2021 Human for the genes we could predict significantly (P* < 0.05) in both men and women. For women, there were no significant pathways after FDR correction, however for men we found 19 pathways at P* < 0.05 and 3 at the stricter 0.005 level. The complete pathway set that came up as significant in men (P* < 0.05) is displayed in Figure 5.

**Figure 5:**
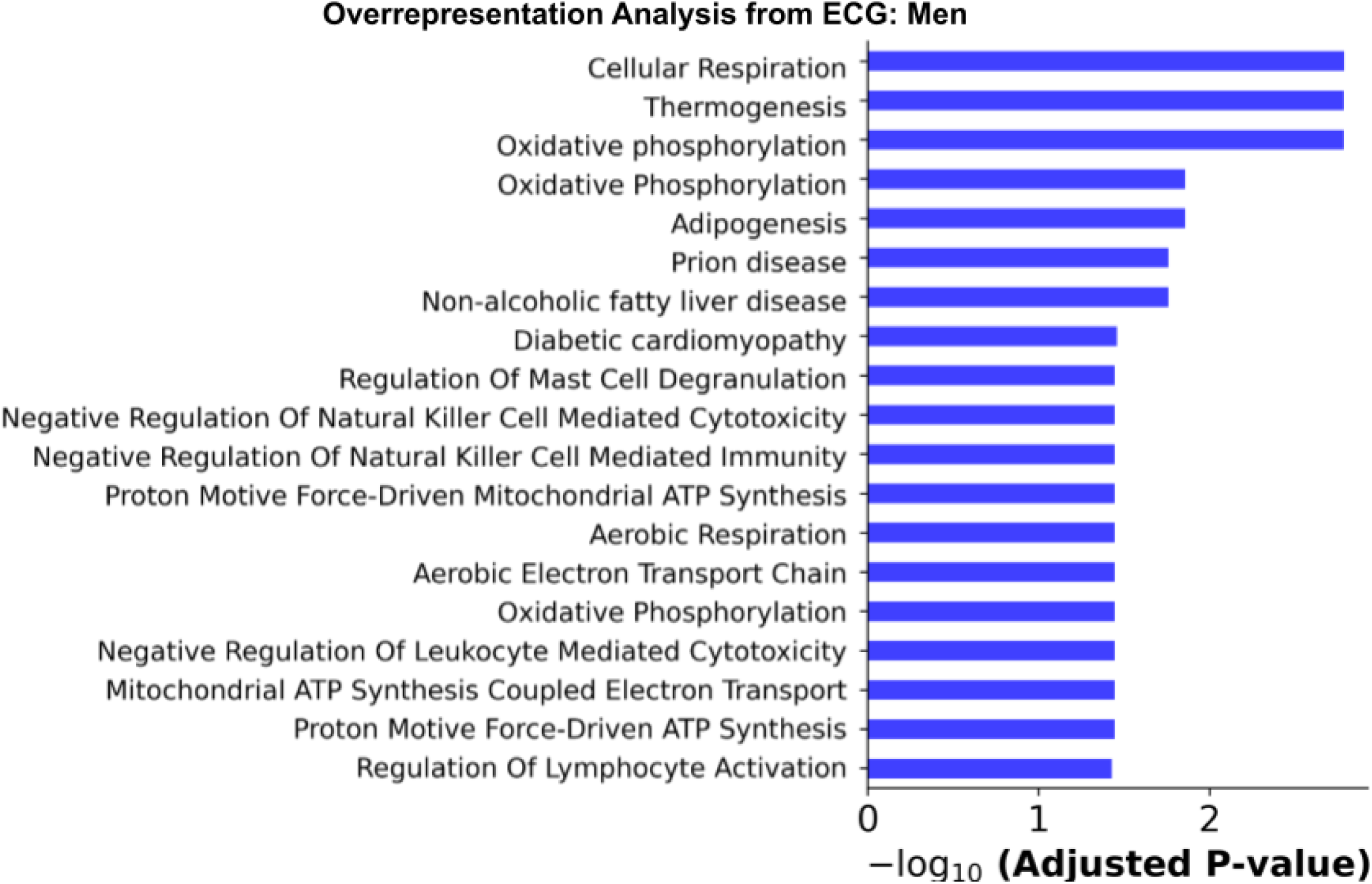
The electron transport chain and immune response are implicated in learned ECG baseline features. Significantly over-represented pathways among genes with well-predicted expression (in men) from ECG embeddings

### The ETC is centrally implicated in Cardiovascular Disease

The most dominant pathway set from our predictions of gene expression is that pertaining to the mitochondrial electron transport chain, in which we identified 9 pathways that were significantly over represented. These are, including some pathways duplicated across multiple gene sets: Cellular Respiration (GO:0045333, P*<0.005), Oxidative Phosphorylation (KEGG:hsa00190, P* < 0.005 & GO:0006119, P* < 0.05) & MSigDB Hallmark, P*<0.05) Proton Motive Force-Driven Force-Driven ATP Synthesis (GO:0015986, P* < 0.05), Mitochondrial ATP Synthesis (GO:0042776, P* < 0.05), Mitochondrial ATP Synthesis Coupled Electron Transport (GO:0042776, P* < 0.05) Aerobic Electron Transport Chain (GO:0019646, P* < 0.05), and Aerobic Respiration (GO:0009060, P* < 0.05). This replicates existing findings on the central importance of mitochondrial function in cardiovascular disease development^18^.

### Embeddings directly predict expression of genes along pathways containing GWAS hits

In our GWAS results, we found a hit in (rs113004948, P < 4e-8) in *CDKL3*. This gene also has a pre-existing known genetic association for Eosinophil count, a known correlate for cardiac complications. From this, we concluded that Delta ECG shares a genetic signal with CVD correlates. However, from the baseline embeddings of patients ECG we could also predict genes in which the pathway for Negative Regulation Of Leukocyte Mediated Cytotoxicity (GO:0001910, P* < 0.05) was over represented. The same can be said for our hit in *MEGF6*, which has known associations for Systemic Mastocytosis^16^, while for RNA in men, the pathway for Mast Cell Degranulation (GO:0043304, P* < 0.05) was significantly over represented in our gene expression predictions. These results indicate that changes in cardiovascular state over time do not just share a genetic basis with eosinophil count and mast cell functioning, but that extracted features from ECG using our model are directly predictive of biomarkers pertaining to the functional role of the two at the cellular level, even in PBMCs.

### Purine Metabolism: Results from an application of enrichment analysis

Lastly, we sought to leverage the RNA expression values directly in our pathway analyses. To do this, we applied gene set enrichment analysis with the case/control label set to a binary discretization of the distance between a patient’s baseline and follow-up appointment (see Methods). Here, women had no pathways, but in men, one pathway came up as significant: Human Purine Metabolism (KEGG:hsa00230, P* < 0.05) whose connection to CVD has been well established previously ^19,20^. This once again can serve as validation of our method.

## Discussion

To understand the genetic basis of individual-level changes in cardiovascular state over time, we conducted genetic analyses on the difference in learned patient cardiovascular state from a novel deep learning model.

There are a few previous works that combine cardiac SSL and genetic studies, though existing studies with ECG are largely cross-sectional, and focus on the factors that cause an between-patient differences in cardiovascular state at a single time point ^1,21^. With respect to repeated ECG recordings over time, only one previous work utilized data (from the UK Biobank) from repeat appointments, however similarity to baseline was used as a quality-control filter as opposed to a learning signal ^22^. Aging is best observed within a single patient over time. Our work is the first to consider the genetic underpinnings of cross-temporal contrastive representations from ECG of the same patient. As such, our representations are perhaps more holistic views of cardiovascular health than those pertaining to a single disease outcome. More generally, our framework can be applied to any medical modality to quantify and understand the genetic underpinnings of temporal changes in phenotypic state as assessed by medical tests.

Still, our work is not without limitations. We used generalized patient identification metrics for each task separately to verify that our model was correctly meaningful patient-specific representation from ECGs. Our “intra appointment” performance was lower than “inter “tasks. This is expected: periodicity within ECG signals across time means that predicting the first half from the second half of an ECG signal is much easier than predicting what a recording a few years down the line and with slight differences in electrode placement in different measurement occasions will look like. Still, our performance on both embedding metrics is indicative of the powerful representations captured by our model.

As well, the sex-based differences we found in the over-representation and enrichment analyses are a result of the poorer gene expression predictions in women. This may be a result of the well-established lower CVD prevalence in women. However, we also cannot exclude the possibility that this has occurred as a result of batch effects or other problems with our ECG recordings, RNASequencing experiment, or both.

The majority of the pooled HPP cohort studied here was relatively healthy. More significant cardiovascular deterioration, and therefore larger Delta ECG would be expected in a cohort comprised of more individuals with CVD. Still, the fact that among largely healthy individuals we were still able to detect disease signals is further proof of the strength of our method.

Numerous works have shown the dependency of contrastive learning methods on using large datasets, and thus potential results are always a function of cohort size. Transfer effects across different datasets within physiological deep learning have been found to be highly significant across datasets^23,24^ and thus we trained our model on our dataset directly, as opposed to training on a larger collection of ECGs such as the UK Biobank. Several future directions are suggested for our work, the first of which is training on a larger sample size. This would not only benefit our SSL: our sample size was below 5000, which is the minimum at which traditional heritability estimation methods, i.e (LD-based estimation and GREML) are sufficiently powered. Fitting the model on a larger cohort would allow us to explore the heritability of Delta ECG, and the genetic correlation between Delta ECG and other traditional cardiac phenotypes. As well, exploring more than one time point per patient could enable deeper exploration of patient trajectories.

## Methods

### Code and Model Weights Availability

All model weights and associated code have been deposited on GitHub, accessible here.

### Architecture Specifications

We aimed to build a novel architecture within which to evaluate our results. The natural starting place in the field of ECG analysis is the one dimensional convolutional neural network (1d-CNN). To improve expressibility as opposed to simply using CNNs, our architecture additionally utilizes transformer encoder layers. Combining CNNs with transformers allowed us to enhance ECG representation learning by using the CNN to generate a key set of temporal tokens, while the self-attention layers were able to model the deep global relationships between them.

As we had two windows for each appointment, we averaged their embeddings to arrive at a single vector per visit, and applied cosine similarity to these vectors.

### Training Details

We initialized the model described in Table S1 (See Supplement) with random weights, and trained using the AdamW optimizer^25^, with an initial learning rate of 1e-5, which we found empirically gave the best results on the validation set. We reduced the learning rate by a factor of 5 for every 5 epochs that the validation loss did not decrease, and after 15 epochs of non-decreasing validation loss, we stopped training the model, and checkpointed the weights for later use. As is standard in the contrastive learning literature, we used the linear projection head for training only, and dropped the projection during inference (generating embeddings per-person). We fit a residual convolutional tokenizer following a standard architecture scheme^26^, and followed by 12 transformer encoder layers^27^ using the BERT Base hyperparameters^28^, including 12 attention heads, a hidden dimension of 3,024. In each forward pass, we pass each recording through the 1d-CNN encoder, yielding a sequence of 58 tokens aligned in time, over 768 channels. We set 768 to be the embedding dimension of the transformer network, and add positional embeddings. We then add a 59th additional token to the model whose value is instantiated randomly (Gaussian), similarly in essence to the CLS token of a standard Vision Transformer (ViT)^29^, which interacts with all other tokens and whose value is sent to the linear projection before becoming the final embedding. For each window per person we obtained a 768 dimensional embedding vector from our model. We used Python Version 3.11.3 for all analyses. We fit all models using PyTorch version ‘2.3.1+cu121’^30^ with the Nvidia Quadro RTX 8000 Graphics Processing Unit (GPU).

### ECG Dataset

We began with a dataset comprising 16,775 ECGs, with 11,933 recordings from baseline visits, and the remainder from repeat appointments. Using 12 Lead NORAV ECG machine –PC-ECG 1200^31^ with an integrated electrodes chest belt for precordial leads. The ECGs were all sampled at a rate of 1000 hz, for 10 seconds minimum, before applying a 50Hz AC noise filter and an EMG muscle noise filter at 35 Hz. The baseline filter was set to on, and we used 16 bit resolution. We split each recording into two non overlapping windows 4 seconds in length across all 12 channels. For the SSL pre-training task, we split the dataset randomly 90/5/5 into train/test/validation over patient identity. We excluded automatically-detected changes related to heart rate, artifacts (identified by focal changes in only part of the leads), and lead misplacement.

### Embedding Neighbourhood Metrics

As we had two windows per recording, we averaged the embeddings from both windows to apply patient identification metrics^8^. We could have of course done this without averaging the embeddings from each window (treating each window separately), however we wanted to encourage all windows from both recordings to be close together, as opposed to just one of each. Within the sets of first and second appointments, we too can apply this metric without averaging the two windows: instead seeing whether the first half a patient’s ECG is closest to the second as compared to all other first or second windows.

### PBMCs Isolation

Blood samples were collected with BD Vacutainer® CPT™ Cell Preparation Tube with Sodium Citrate and Ficoll (BD Ref# 362760), and PBCMs were purified according to the manufacturer’s instructions with minor modifications. Briefly, the PBMCs were collected from the CPT tubes after 20-minute centrifugation at 22oC followed by two washing steps with Washing media (0.9%RPMI 1640, Thermo Fisher Scientific, and 0.1%FBS, Sigma-Aldrich). The cells were resuspended with resuspension solution (0.5% RPMI 1640, Thermo Fisher Scientific and 0.5%FBS, Sigma-Aldrich) and frozen at −80oC until processing. All processes were performed on a Tecan Evo 100 automated platform.

### RNA extraction and library prep and sequencing

RNA was extracted from frozen PBMCs with an All-prep DNA/RNA 96 (4) kit (QIAGEN, Cat# 20-80311) on a Tecan Evo 200 automated platform. Libraries for bulk mRNA sequencing were prepared with mcSCRB-seq methodology^32^. All steps were identical except for the final step of tagmentation. 96 amplified cDNA samples were pooled and 12ng from each pool was tagmented by adding 1uL of TDE1 and 2x Tagment DNA buffer (Illumina, Ref# 20034197) in 60uL followed by PCR amplification with Kapa HiFi HotStart ReadyMix (Kapa Biosystems) and 5 μM IDT for Illumina Nextera DNA Dual Indexes. The PCR conditions were: 3’ at 72°C, 30’’ at 95°C followed by 14 cycles of 10’’ at 95°C, 30’’ at 55°C, 1’ at 72°C and final elongation for 5’ at 72°C. PCR clean-up and size selection was performed using SPRI beads for each pool and eluted in 12uL. Libraries were sequenced to a minimum depth of 5M reads per sample on a NovaSeq 6000 instrument with a NovaSeq S1 v1.5 100-cycle kit (Cat# 20012865; Illumina).

Deduplicated count data for the gene expression was computed using a previously validated pipeline^33^. These were then converted to counts per million mapped reads through normalization.

### RNA Expression Prediction

For downstream task prediction, we stored embeddings from each person into a tabular dataset, and fit Ridge linear regression models over them to predict the output. We used cross-fold-validation over the dataset to predict the test prediction performance.

### Enrichment Analysis

We discretized the Delta ECG distribution by assigning patients to the “case” category if their distance between their matched appointments was above the mean of the matched distance distribution (0.9), and “control” group otherwise. Using GSEAPY^34^, we ran over gene set enrichment analysis on all 3000 genes for both men and women, ranked by corrected prediction P value from ECG embeddings. We used 5000 permutations, and the log (base 2) ratio of classes.

### Phenotypes

We set the data collection interval to be from the commencement of the HPP, in January of 2019 to June 2024.

### RNSeq: Ranked Multiple Hypothesis Testing Correction

Our RNASequencing dataset includes expression values for over 10k genes per patient, though the data itself is sparse. The remaining genes which are expressed across people could be picked based on a combination of higher variance, higher expression, or randomly. Either way, a set number of genes needed to be subsampled for the experiment. As multiple hypothesis correction is a function of the number of tests performed, to exclude the possibility of hand-picking this number of genes so that our results were significant, we employed sequential multiple hypothesis correction in the following way: we first ordered all genes based on decreasing population-level variance. For the first (most-variable) gene, we left its P-value intact. For the nth-gene in the list, we adjusted its P value using Bonferonni adjustment for n tests. This is essentially an analogue of the Holm-Bonferonni method, without ordering the P values from lowest to highest first. Results from experiment consisting of the top n genes would therefore only be penalized for those genes.

### Genome Wide Association Study

Our methods for GWAS and LD clumping have been described previously^9^. We use the same methodology for our eQTL analyses, replacing the clinical phenotypes with the normalized expression values for each of the 66 candidate genes across the population.

## Acknowledgements

We thank members of the Segal lab for useful discussions. E.S. is supported by the Crown Human Genome Center; Larson Charitable Foundation New Scientist Fund; Else Kroener Fresenius Foundation; White Rose International Foundation; Ben B. and Joyce E. Eisenberg Foundation; Nissenbaum Family; Marcos Pinheiro de Andrade and Vanessa Buchheim; Lady Michelle Michels; Aliza Moussaieff; and grants funded by the Minerva foundation with funding from the Federal German Ministry for Education and Research and by the European Research Council and the Israel Science Foundation.

## Author Contributions

Z.L. conceived the project, designed and trained the neural network, performed the genetic analyses, interpreted the results, and wrote the manuscript. G.L. contributed insights and assisted with the suggestion of ideas for the project. A.G. coordinated and provided the DNA data, coordinated, conducted, and designed the RNA Sequencing experiment, interpreted the results, and wrote the manuscript. A.W. coordinated, conducted, and designed the RNA Sequencing experiment, interpreted the results, and wrote the manuscript. M.P. coordinated, conducted, and designed the RNA Sequencing experiment and wrote the manuscript. Y.T.B and Y.R. interpreted the results, wrote the manuscript, and provided valuable clinical insights. H.R. provided data, interpreted the results, and wrote the manuscript. E.S. conceived, directed, and supervised the project and analyses

## Ethics

The Weizmann Institute of Science review board (IRB) approved the study and its protocols. All identifying details of the participants were erased prior to statistical analysis, so informed consent was waived by the IRB. All participants had full knowledge of data handling, storage, and sharing methods. This information was given to all participants, and is in agreement with the data privacy and protection policy of the Weizmann Institute of Science (https://www.weizmann.ac.il/pages/privacy-policy).

